# Germplasm stability in zebrafish requires maternal Tdrd6a and Tdrd6c

**DOI:** 10.1101/2024.10.04.615777

**Authors:** Alessandro Consorte, Yasmin El Sherif, Fridolin Kielisch, Nadine Wittkopp, René F. Ketting

## Abstract

Germ cell specification is driven, in many species, by germplasm: a collection of phase-separation-based structures in the embryo, formed by RNA and proteins that derive from the cytoplasm of the oocytes. How the formation of germplasm is regulated, especially in vertebrates, remains unclear. In this study, we show that two multi-Tudor proteins, Tdrd6a and Tdrd6c, together are necessary for the stability of germplasm in zebrafish, while the related Balbiani body in the oocyte is largely unaffected. Combined lack of maternal Tdrd6a and Tdrd6c still allows the initial germplasm formation but causes its dispersal during the first hours of development. This results in the absence of primordial germ cells during later development and in sterility of the resulting adult animals. Furthermore, our study suggests that the Prion-like domain of Tdrd6c is relevant for Tdrd6c self-interaction as well as for its interaction with Bucky ball, the organizer of germplasm in zebrafish and that these dynamics are modulated by Tdrd6c Tudor domains. The identification of Tdrd6a and Tdrd6c as required for germplasm stability is an important step in our understanding of how this phase-separated structure is controlled during development.

**Author Summary:** We show that maternal Tdrd6a and Tdrd6c proteins interact with Bucky Ball and are essential for embryonic germplasm stability, germ cell specification and fertility in zebrafish.

## INTRODUCTION

The germline of a multicellular organism consists of a specific set of cells, the germ cells, which are essential for the fertility and the reproduction of the individual. Only these cells have the capability to form the zygote, a totipotent cell from which a whole organism can develop. Two major strategies have been identified for specifying the germ cells: induction and pre-formation (Extavour, 2003). The induction mode relies on signaling pathways, which act between cells of the developing embryo, to trigger a small group of cells to differentiate into so-called primordial germ cells (PGCs). Pre-formation, instead, functions via the deposition of determinants (proteins and RNAs) expressed in the oocytes. Upon fertilization, these maternal determinants specifically enrich within a specific set of cells that will then take on PGC fate.

Although pre-formation mechanisms can differ between species, they all rely on phase-separation-related mechanisms, resulting in the formation of so-called germplasm (Boke et al., 2016; Bose et al., 2022; Brangwynne et al., 2009). In this study we focus on germplasm formation in the zebrafish (*Danio rerio)*. In zebrafish oocytes, the protein Bucky ball (Buc) triggers the formation of a large amyloid-like structure, called the Balbiani body (Bb). This body lies at the vegetal side of the oocyte and enriches many determinants relevant for PGC specification (Boke et al., 2016; Bontems et al., 2009; Marlow & Mullins, 2008). With the growth and maturation of the oocyte, the Bb disperses into small patches at the vegetal cortex of the cell. After fertilization, the Bb components migrate to the animal pole, where the embryo develops, and organize into relatively small cytoplasmic droplets. With the start of the first two cell divisions, many of these small condensates align along the forming cleavage furrows of dividing cells. This process results in the formation of four clusters of droplets: these are referred to as germplasm. By the onset of gastrulation, the germplasm ends up in four groups of cells, driving their development as PGCs (Raz, 2003). Although we know much about the timed assembly and distribution of germplasm, many proteins regulating these transitions remain unknown.

Buc is a known organizer of the Bb in zebrafish oocytes (Bontems et al., 2009; Dosch et al., 2004; Marlow & Mullins, 2008). Interestingly, Buc is a largely intrinsically disordered protein (IDP) with a predicted Prion-like domain (PrLD) at its N-terminus. This PrLD has been shown to be necessary to confer amyloid-like features to the Bb (Boke et al., 2016). This is also consistent with the observation of other phase-separation related studies where IDPs and PrLDs have been found to be relevant for the regulation of different condensates (Alberti et al., 2009; Banani et al., 2017; Kroschwald et al., 2015; Malinovska et al., 2013). The Bb, however, remains one of the few structures known thus far to show clear amyloid characteristics, while maintaining a dynamic nature: it disperses and reassembles.

In previous work, we have shown how a multi-Tudor domain containing protein, Tdrd6a, interacts with Buc and affects its mobility (Roovers et al., 2018). The absence of Tdrd6a protein during zebrafish early development causes reduction in the number of PGCs at the larval stages, accompanied by mild germplasm defects. Analysis of the Buc-Tdrd6a interaction showed it was mediated through binding of methylated arginine/glycine (RG) motifs in Buc, presumably by the extended Tudor Domains (eTDs) of Tdrd6a. This study also revealed the presence of Tdrd6c, a paralog of Tdrd6a, which was suggested to also bind Buc, again likely via its eTDs (Roovers et al., 2018).

In this work we show that Tdrd6a and Tdrd6c both contribute to the stability of the embryonic germplasm, and that they act redundantly to ensure PGC formation and fertility of the zebrafish. Furthermore, we demonstrate that the Buc-Tdrd6c interaction is not mediated by its Tudor domains but the PrLD of Tdrd6c, which is not found in Tdrd6a. Thus, our work provides deeper insights into the regulation of germ cell pre-formation mechanisms and describes a first mutant line of zebrafish (*tdrd6a/c* double mutants) in which the embryonic germplasm is fully destabilized during early development, without affecting embryonic viability and growth of the animal.

## RESULTS

### Tdrd6c has seven extended Tudor domains and a Prion-like domain

In zebrafish, three Tdrd6 paralogs are encoded within the genome: *tdrd6a* (ENSDARG00000070052) and *tdrd6b* (ENSDARG00000014039) on chromosome 20 and *tdrd6c* on chromosome 17 (ENSDARG00000089954). Despite their localization on different chromosomes, the structure of the loci of *tdrd6b* and *tdrd6c* appears similar, with three exons contributing to their open reading frames (ORFs) in contrast to *tdrd6a* which displays 15 coding exons (Fig. S1A). Interestingly, all *tdrd6* loci of zebrafish share a large exon of over 5000 base pairs (bp), encoding the majority of the final protein product, while all the other coding exons are below 200 bp in size. Furthermore, all *tdrd6* genes code for proteins of similar size and structure, harboring seven predicted eTDs distributed in their protein sequences. However, *tdrd6b* has not been reported to be expressed in the zebrafish germline and was not identified as an interactor of germplasm components in mass-spectrometry analyses of different protein pull-downs (Y. Liu et al., 2022; Roovers et al., 2018). For these reasons, we focused on *tdrd6a* and the uncharacterized *tdrd6c*.

*D. rerio tdrd6c* encodes a protein of 1814 amino acids in length that contains seven eTDs (Fig. 1A). When visualized via Alpha Fold predictions, the eTDs of Tdrd6c have the additional α-helices and β-sheets typical of eTDs (Fig. S1B) (Jumper et al., 2021; H. Liu et al., 2010; K. Liu et al., 2010). This class of Tudor domains has often been found in the interaction with substrates containing methylated Arginines. In eTDs, four residues form an aromatic cage within the β-barrel core structure, where one Asparagine residue exhibits affinity for methylated-substrates (H. Liu et al., 2010; K. Liu et al., 2010). These are indeed found in the four C-terminal eTDs of Tdrd6c (Fig. S1C). However, the first three eTDs of Tdrd6c do neither contain the aromatic cage, nor do they have the conserved Asparagine, suggesting that these N-terminal eTDs of Tdrd6c could exhibit different substrate recognition (Fig. S1C). This organization is similar to that of Tdrd6a (Fig. S2), although the number of eTDs with and without an intact aromatic cage differs between the two proteins. Based on sequence homologies it seems likely that specific eTDs duplicated or disappeared when Tdrd6a and Tdrd6c arose from a shared ancestral gene (Fig. S2A,C)

**Fig. 1.**
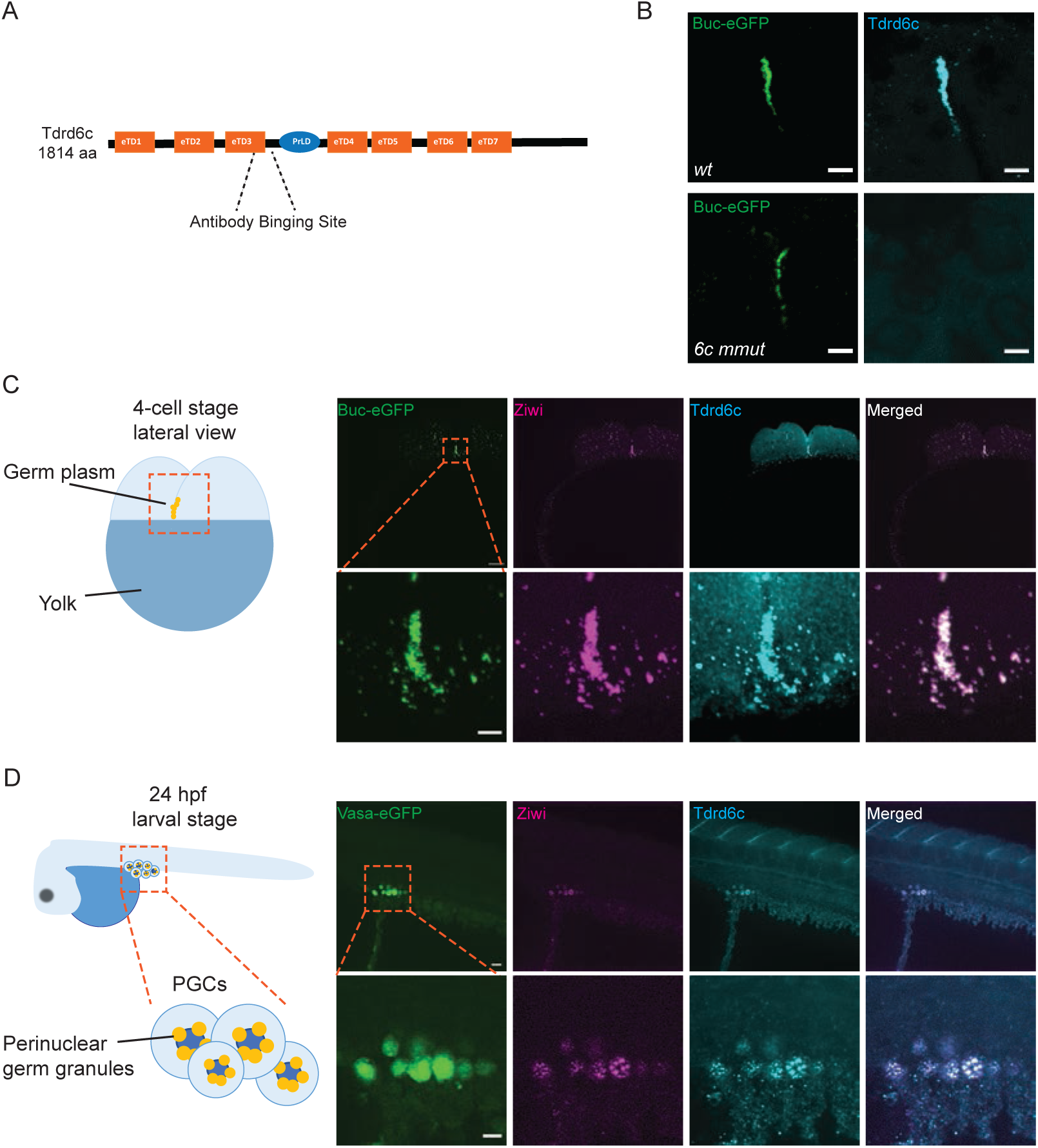
Tdrd6c localization in zebrafish embryos. (A) Scheme of Tdrd6c protein structure; orange boxes indicating Tudor domains (eTDs) and the blue ellipse the predicted prion-like domain (PrLD). (B) Validation of the specificity of the Tdrd6c-antibody, showing its signal co-localizating with Buc-eGFP in *wt* embryos (top) but being absent in embryos collected from *tdrd6c^-/-^* mutant mothers. (C) On the left, a scheme of the lateral view of the 4-cell stage zebrafish embryo, with yellow Gp droplets distributing on the cleavage furrow; on the right, the panel showing co-localization of BucGFP (green), Ziwi (magenta) and Tdrd6c (cyan); below are zoom in images. (D) On the left, a scheme of the 24 hpf zebrafish larva, where PGCs are localized ventrally in proximity of the yolk (dark blue); yellow circles indicate perinuclear granules; on the right, the panel showing VasaGFP-positive PGCs, whith co-localization of Tdrd6c (magenta) and Ziwi (cyan) into perinuclear granules; below is the zoom in panel of the PGCs. Orange dashed squares indicate the zoom in areas; scale bars: 20 µm for (B), 50 µm for (C), 10 µm for (D) and 5 µm for zoom in of (C) and (D).

The presence of a predicted Prion-like domain (PrLD) of 92 amino acids between the third and fourth eTD domains in Tdrd6c distinguishes it from Tdrd6a (Fig. S3). PrLDs are known to be involved in the regulation of phase-separation processes (Alberti et al., 2009; Shorter & Lindquist, 2005) and this region in Tdrd6c might be relevant in the regulation of germplasm formation during zebrafish development.

### Tdrd6c is maternally inherited and localizes to germplasm and perinuclear granules of PGCs

To investigate Tdrd6c expression and localization we raised an antibody against a unique amino-acid sequence in rabbits and affinity-purified it. Antibody specificity was validated via immuno-staining on embryos derived from *tdrd6c* mutant mothers (see below), an experiment that also provided clear evidence of Tdrd6c localization to zebrafish germplasm during early embryogenesis (Fig. 1B). Indeed, we found that Tdrd6c was present in the germplasm marked by the transgenic line *Tg(buc:buc-eGFP)* and Ziwi antibody-staining at the 4-cell stage (Fig. 1C) (Houwing et al., 2007; Riemer et al., 2015). At 24 hours post fertilization (hpf), we detected Tdrd6c enrichment in PGCs marked by expression of *Tg(vasa:eGFP)* reporter (Fig. 1D) (Krøvel & Olsen, 2002). More specifically, the signal localized to perinuclear granules in PGCs, highlighted by Ziwi (Fig. 1D) (Houwing et al., 2007). Altogether, these results show that Tdrd6c is maternally provided, present in embryonic germplasm of zebrafish and can still be found in PGCs at 24 hpf.

### A transgenic line expressing a Tdrd6c-mKate2 fusion protein marks *nuage* and germplasm

To observe Tdrd6c localization and dynamics in living embryos we generated a transgenic line expressing a Tdrd6c-mKate2 fusion protein using the transposon-based Tol2-transgenesis technology (Kawakami, 2007). The designed transgene (*tg(ziwi:tdrd6c-mKate2-tdrd6c3’UTR)*) combines the *ziwi* promoter, the *tdrd6c* open reading frame (ORF), the mKate2 ORF and the *tdrd6c* 3’UTR (Fig. 2A). The *ziwi* promoter was chosen in order to drive expression during oogenesis (Leu & Draper, 2010) and the mKate2 fluorophore for combination with available GFP-expressing transgenic lines (Shcherbo et al., 2009).

**Fig. 2.**
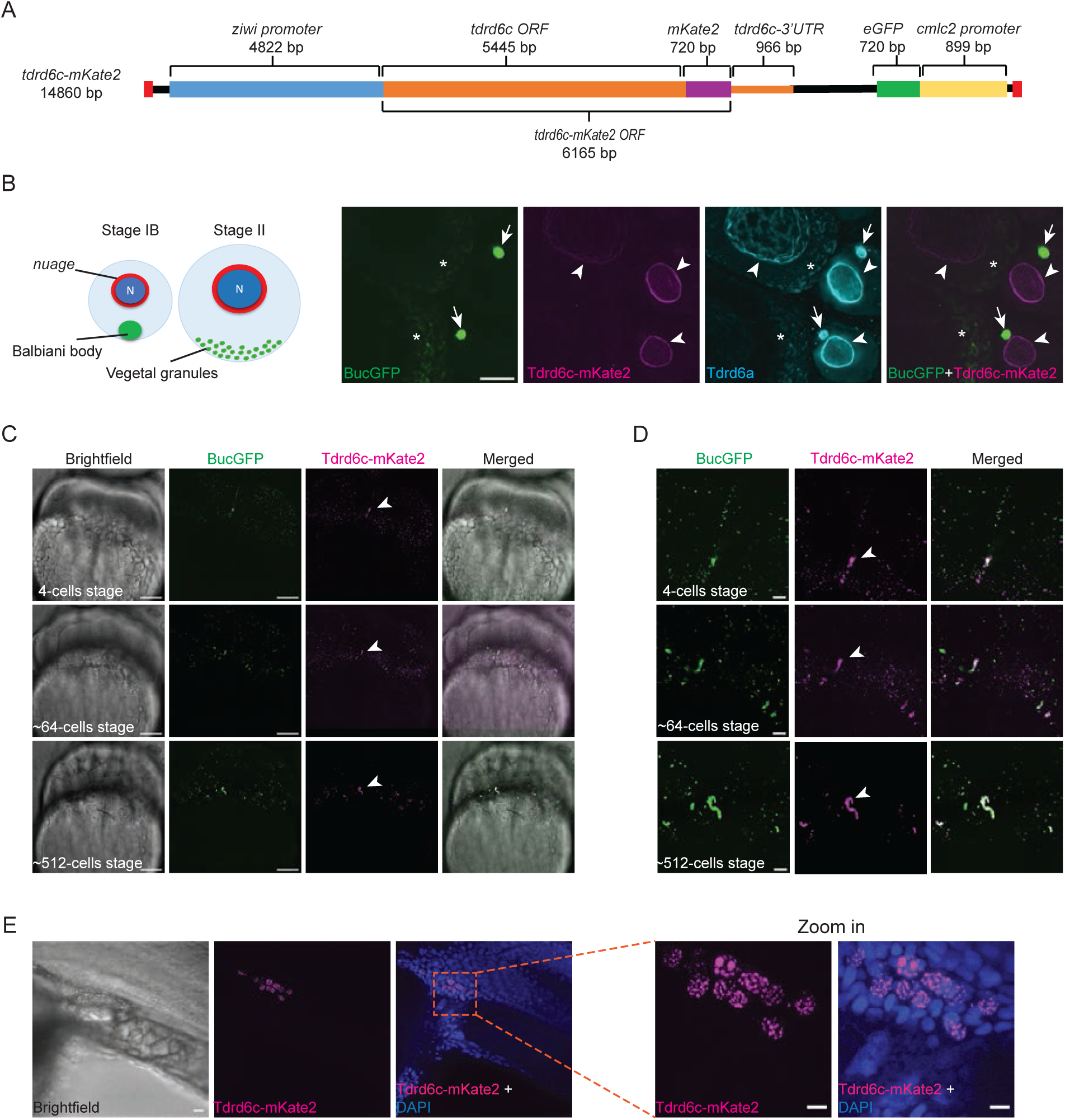
Expression of the Tdrd6c-mKate2 transgene. (A) Schematic representation of the *tdrd6c-mKate2* transgene inserted via Tol2-technology; in red are the Tol2 transposition sites; in black linker sequences. (B) On the left, schematic representation of oocytes stages IB and II (nuclei in dark blue, perinuclear *nuage* in red and Bb and Vg in green); on the right, panel showing BucGFP (green) and Tdrd6c-mKate2 (magenta) expressing oocytes stained with Tdrd6a-antibody (cyan); Tdrd6c-mKate2 localization is restricted to the perinuclear *nuage* (arrowheads) and does not appear enriched in the Bb (arrows) or Vegetal granules (asterisks). (C) Tdrd6c-mKate2 (magenta) localizes to the Gp (arrowheads) together with BucGFP (green), at different stages of embryogenesis. (D) Zoom in of (C). (E) Localization of Tdrd6c in perinuclear granules at 24 hpf (DAPI in blue) with zoom in on the right. Scale bars: 20 µm for (B), (D) and (E), 50 µm for (C) and 5 µm for zoom in of (E).

We detected expression of Tdrd6c-mKate2 in oocytes of different stages, where the signal enriches around the nucleus (nuage) but did not detect enrichment in the Bb (Fig. 2B), which makes it distinct from Tdrd6a (Roovers et al., 2018). After fertilization, maternally provided Tdrd6c-mKate2 localized to germplasm droplets marked by Buc-eGFP and enriched at the cleavage furrows of the dividing blastomeres (Fig. 2C, D and MovieS1). At later stages, the signal of Tdrd6c-mKate2 was detected in cells at the genital ridge of zebrafish larvae, which corresponds to the area where PGCs settle for further development (Fig. 2E). In particular, the Tdrd6c-mKate2 signal was enriched within granules around the PGC nuclei: the perinuclear germ granules. These observations are in agreement with the results collected by antibody-staining and provide further evidence for the localization of Tdrd6c to embryonic germplasm structures and to the nuage of zebrafish oocytes.

### Maternal Tdrd6c affects PGC formation

In order to create a *tdrd6c* mutant allele, we targeted its large exon with CRISPR-Cas9 using two guide-RNAs (gRNAs) (Fig. 3A) (Hwang et al., 2013; Jinek et al., 2012). We successfully generated a *tdrd6c* knock out allele (*tdrd6c^mz69^*) that failed to express Tdrd6c, as its signal was not detectable after fertilization in embryos deriving from homozygous mutant mothers (Fig. 1B). Homozygous mutant *tdrd6c* animals were viable under standard conditions, with fish exhibiting normal development and behavior. Also, these animals did not exhibit a bias towards male or female differentiation and all tested adult fish were able to produce offspring.

**Fig. 3.**
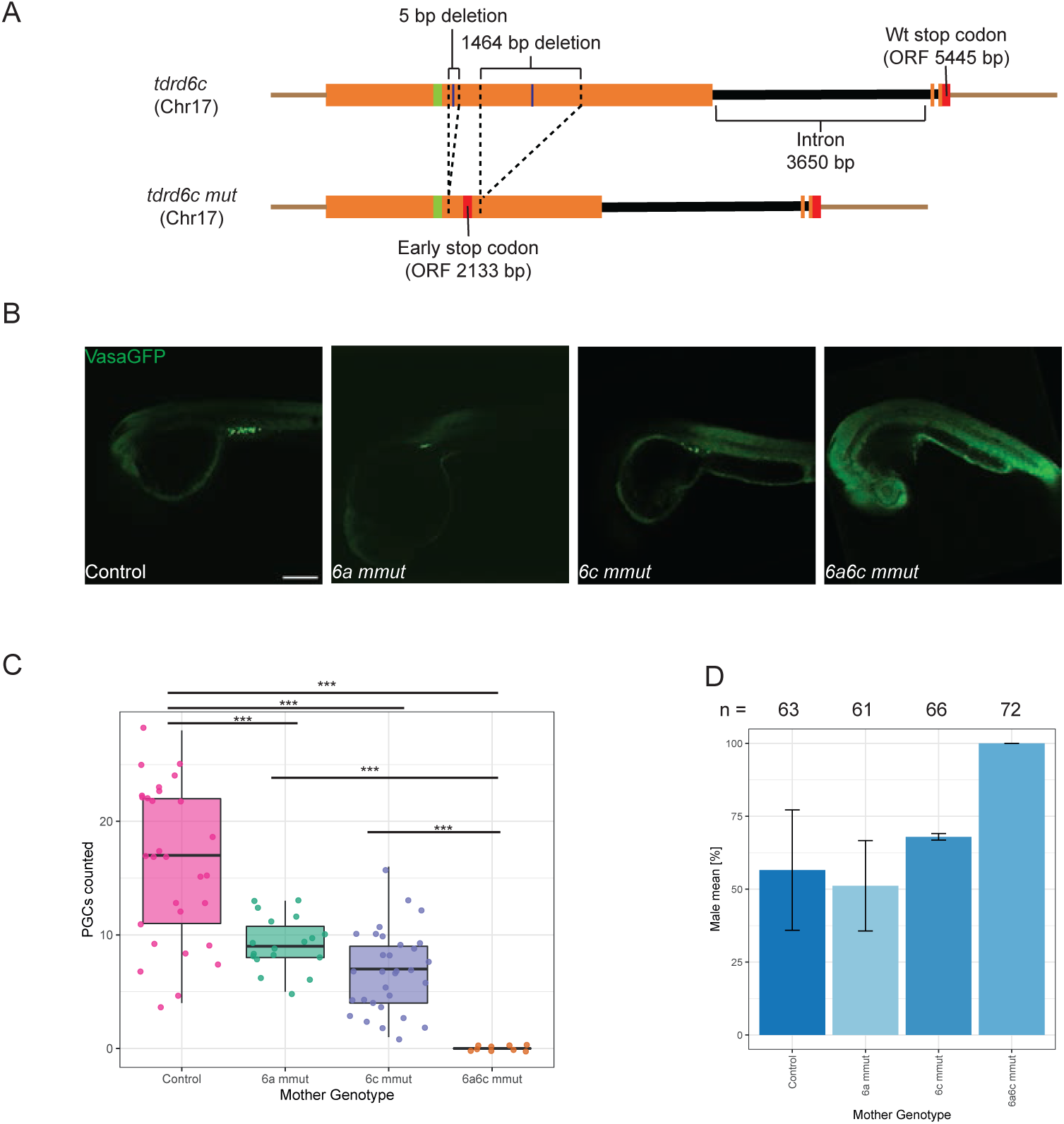
PGCs defects in offspring from *tdrd6a/c* mutants. (A) Schematic representation of *tdrd6c* knock out allele. Black bars are intronic regions, orange boxes represent exons, red boxes stop codons, light green box the epitope recognized by Tdrd6c-antibody and blue lines are targets of the 2 injected CRISPR-RNAs. (B) VasaGFP-positive PGCs in embryos from mothers of different genotypes, showing reduction in *6a mmut* and *6c mmut* and total absence in *6a6c mmut* (scale bar = 50 µm). (C) Quantification of PGC counts in embryos 1 dpf. Significance: (***) = p-value < 0.005. (D) Percentage of males among offspring from females of the indicate genotypes (we never obtained any female offspring from 6a6c double mutant females) (n refers to number of fish that reached 3 months of age for each population). In both (C) and (D) error bars represent standard deviation.

To investigate if PGC development was affected by the lack of maternally provided Tdrd6c protein, we combined the *tdrd6c^mz69^* allele with *Tg(vasa:eGFP)*, which is expressed at all stages of germ cell development. We found that embryos from *tdrd6c^-/-^* females, which we refer to as *tdrd6c* maternal mutants (*6c mmut*), had reduced numbers of PGCs at 24hpf, compared to wild-type (Fig. 3B, C). This phenotype is similar to what was observed in *tdrd6a* maternal mutant embryos *(6a mmut)* (Roovers et al., 2018). We conclude that maternally provided Tdrd6c is involved in PGC formation, but is not essential for this process.

### Simultaneous absence of maternal Tdrd6a and Tdrd6c disrupts PGC formation

Next, we bred animals carrying mutations for both *tdrd6a* and *tdrd6c* and characterized their phenotypes. Like for the single mutants, *tdrd6a^-/-^; tdrd6c^-/-^* double mutants obtained from heterozygous parents developed normally and were fertile. Strikingly, however, double mutant *6a6c mmut* embryos showed complete absence of PGCs at 24 hpf (Fig. 3B, C). This indicates that the maternal inheritance of at least one of the Tdrd6 proteins is crucial for the specification of PGCs.

Next, we raised *6a6c mmut* fish to adulthood and compared their growth and maturation to wild type control populations and to *6a mmut* or *6c mmut* fish. These were collected on the same day and reared at the same density, trying to minimize any stochastic influence from the environment to the different batches. After three months, all populations looked mature, with no superficial phenotype. However, when assessing sex ratios in each tank, we did not find any female fish in the *6a6c mmut* adult populations (Fig. 3D). Furthermore, 31 *6a6c mmut* adult males, obtained from different crosses, were tested for fertility and were not able to fertilize eggs upon mating with different *wt* females (Fig. S4A). The dissection of these males showed that *6a6c mmut* adult fish had severely underdeveloped gonads (Fig. S4B, C). This is consistent with previous studies that found that PGC loss leads to sterility and a strong bias towards male differentiation in zebrafish (D’Orazio et al., 2021; Gross-Thebing et al., 2017; Houwing et al., 2007; Siegfried & Nüsslein-Volhard, 2008; Weidinger et al., 2003). From these observations, we conclude that maternal Tdrd6a and Tdrd6c act redundantly in the process of PGC specification and that their simultaneous absence prevents the formation of germ cells during early development, resulting in sterility.

### Maternal inheritance of Tdrd6a or Tdrd6c is crucial for germplasm stability and enrichment during cell divisions

We next employed the Buc-eGPF expressing transgene to test the effects of maternal Tdrd6a and Tdrd6c on germplasm in the early embryo. (Riemer et al., 2015). Upon analysis of germplasm droplet abundance and volumes in *6a mmut* and *6c mmut* embryos we did not observe any significant differences in comparison to control populations (Fig. 4A, B). However, in *6a6c mmut* embryos we observed that the Buc granules completely disappeared around 3 hpf (Fig. 4A, B). This loss of Buc granules appears to be a gradual process, as can be appreciated from tracking granules over time (MovieS2 and MovieS3). We conclude that in absence of both maternal Tdrd6a and Tdrd6c germplasm initially forms, but that it is instable and gradually lost during the first hours of development.

**Fig. 4.**
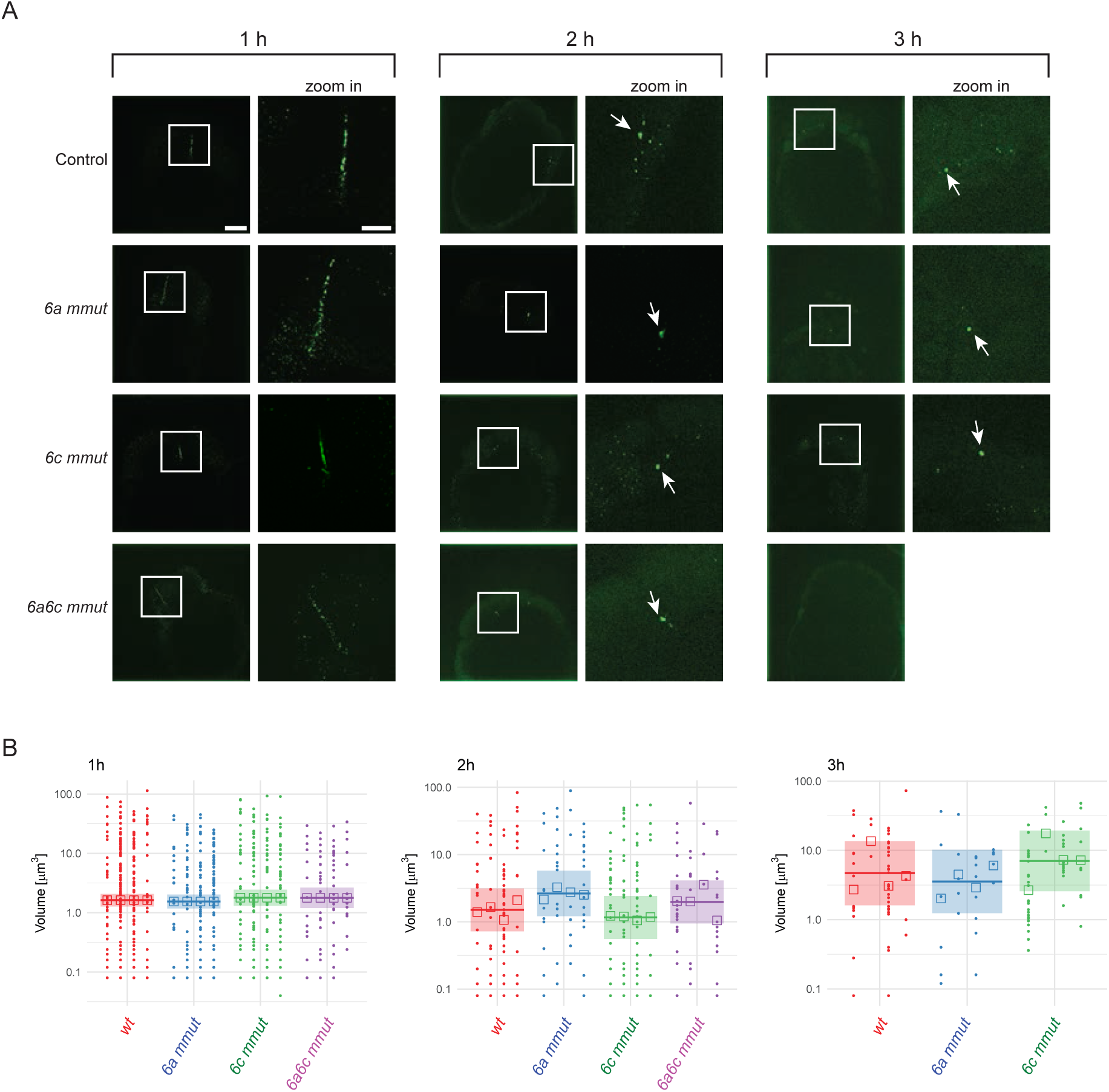
Germplasm defects in *6a6c mmut* embryos. (A) Germplasm, marked by BucEGFP, at different stages of embryogenesis embryos from mothers of indicated genotypes; different stages are listed from left to right as 1 hour (h), 2h and 3h; genotypes (maternal mutants) are reported on the left, from top to bottom; squares indicate the area of zoom in; arrows highlight germplasm condensates; scale bars = 50 µm and 20 µm for zoom in. (B) Quantification of BucGFP signal. Both volume of individual Buc granules, as well as their numbers were quantified in embryos of the indicated genotypes and at the indicated time points. For each indicated genotype, 4 individual embryos were measured (represented by each different column of dots per group). Shaded bands highlight 95% confidence intervals for the mean volume. Analysis in log-scale without multiple testing correction. The graph at 3 h (on the right) does not report measurements for *6a6c mmut* as no germplasm condensates were found in those individuals.

### The Tdrd6c PrLD triggers protein interaction and self-assembly in BmN4 cells

Next, we aimed to study the aggregation behavior of Tdrd6c in more detail. For this, we chose to express different protein variants as N-terminally tagged mCherry fusion proteins in *BmN4* cells (Fig. 5A) (Shaner et al., 2004). We chose these *Bombyx mori*-derived cells because of their germline character and because they are normally cultured at 28°C, which is the same temperature preferred for zebrafish breeding, making this system suitable to test and analyze zebrafish proteins (Roovers et al., 2018). The designed constructs reached similar levels of expression in the cytoplasm of BmN4 cells, as measured by the intensity of the mCherry fluorophore (Fig. 5B).

**Fig. 5.**
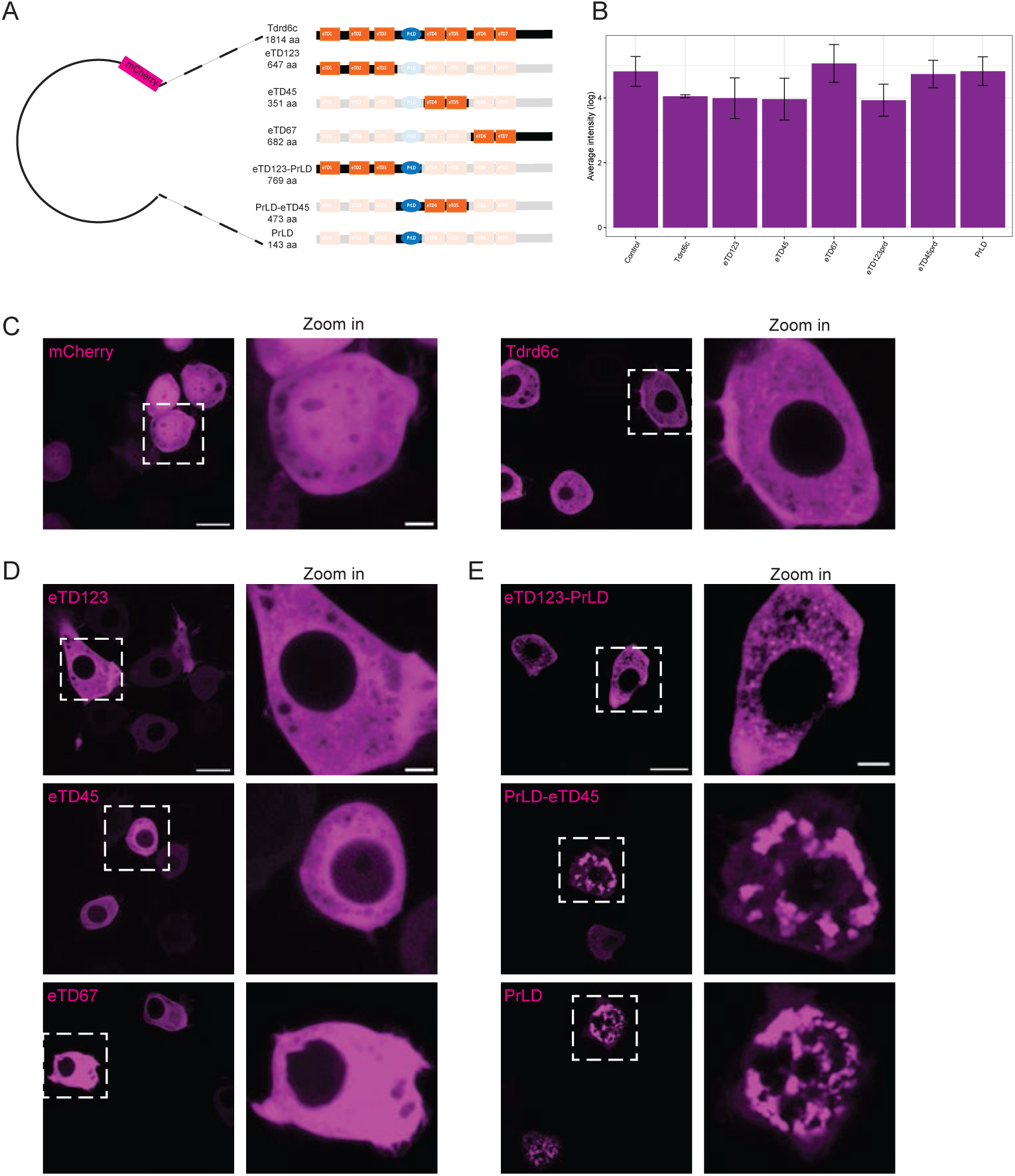
Expression of Tdrd6c constructs in BmN4 cells. (A) Schematic representation of plasmids expressing Tdrd6c constructs for BmN4 transfection. (B) Levels of expression measured for each tagged-Tdrd6c construct based on the total fluorescence signal detected (for each construct 9 cells were measured). Error bars represent standard deviation. (C) Expression of mCherry compared to expression of mCherry-Tdrd6c. (D) Expression of Tdrd6c constructs that diffuse in the cytoplasm of BmN4 cells. (E) Expression of Tdrd6c constructs that exhibit granular expression. Scale bars: 20 µm (C and D) and 5 µm (zoom in of C and D); dashed squares indicate the areas of zoom in.

We started our analysis with transfection of the Tdrd6c full-length protein and observed that its signal was diffuse within the cytoplasm; no aggregation or particular foci of enrichment were observed for full-length Tdrd6c (Fig. 5C). Its exclusion from the nucleus, in contrast to the mCherry control, likely reflects the size of the fusion protein. Constructs expressing combinations of solely eTDs behaved similar to the full-length protein (Fig. 5D). In contrast, expression of the PrLD alone or in combination with adjacent sequences triggered different dynamics. When we expressed mCherry-PrLD or mCherry-PrLD-eTD45, we observed large cytoplasmic aggregates of irregular shape (Fig. 5E). On the other hand, expression of mCherry-eTD123-PrLD exhibited formation of smaller granules (Fig. 5E). These results indicate that the PrLD of Tdrd6c is both sufficient and necessary to confer self-interaction capabilities to Tdrd6c peptides in BmN4 cells. Interestingly, the three N-terminal eTDs, having incomplete aromatic cages, can modulate this self-aggregation behavior while eTD4 and eTD5, which have intact aromatic cages, cannot.

### Tdrd6c PrLD mediates Buc interaction

Next, using this heterologous expression system, we wanted to test a possible interaction between Tdrd6c and Buc. In this system, Buc alone forms droplets that appear spherical and are freely moving and fusing within the cytoplasm (MovieS4) (Roovers et al., 2018). Co-expression of Buc and full length Tdrd6c resulted in occasional co-localization of the two proteins: in the majority of cells that expressed both constructs, Buc was able to form round granules while Tdrd6c was dispersed in the cytoplasm (Fig. 6A). However, we observed a population of cells in which Tdrd6c was found to colocalize with Buc granules (Fig. 6A, B). Furthermore, when expressing different combinations of eTDs of Tdrd6c we did not detect any apparent interaction with Buc, as the latter formed round condensates while eTDs were always found diffuse in the cytoplasm (Fig. 6A). On the other hand, expression of constructs containing the PrLD showed interesting effects. First, the TD123-PrLD protein always co-localized with Buc-positive round condensates in the cytoplasm (Fig. 6A, B). Second, mCherry-PrLD-TD45 formed large aggregates as before, and Buc droplets formed very close to, possibly on the surface of these aggregates (Fig. 6). Interestingly, Buc droplets were immobilized on the PrLD aggregates, in contrast to the Buc granules in absence of the Tdrd6c PrLD (MovieS4, MovieS5, and Fig. S5). These results indicate that the PrLD of Tdrd6c interacts with Buc, and that the Tdrd6c eTDs 1-3 modulate the outcome of this interaction, likely by affecting the aggregation behavior of the Tdrd6c-PrLD itself. Strikingly, the eTDs of Tdrd6c themselves did not display interactions with Buc. These effects are different from what was observed for Tdrd6a, indicating that despite their close homology, Tdrd6a and Tdrd6c interact with Buc in different ways.

**Fig. 6.**
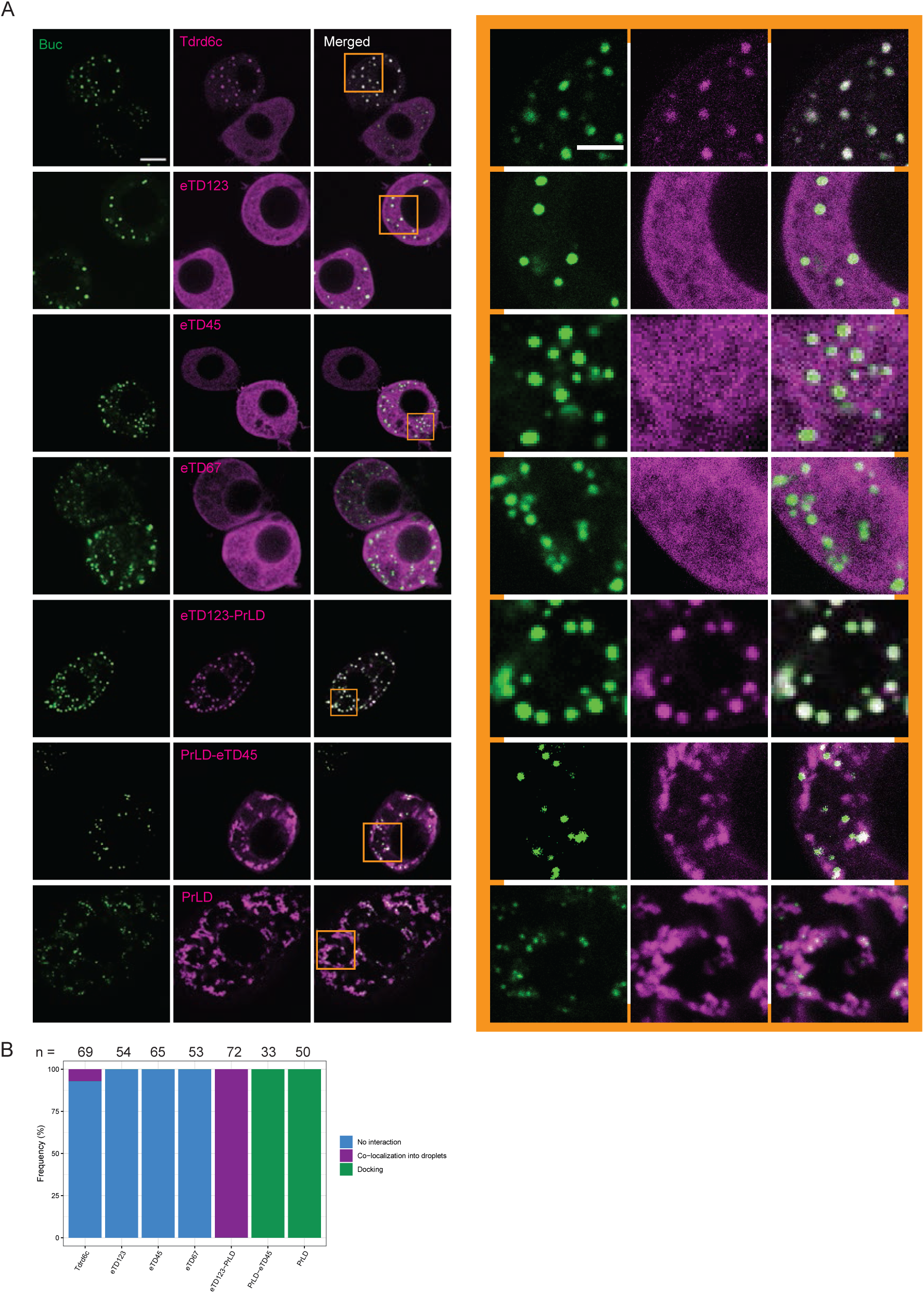
Co-expression of Tdrd6c constructs together with Buc in BmN4 cells. (A) Co-expression of BucGFP and different mCherry-tagged variants of Tdrd6c eTDs (scale bar = 10 µm); on the right, the zoom in of the area highlighted by orange squares (scale bar = 5 µm) (B) Quantification of the frequency observed of the different kind of interactions of mCherry-tagged Tdrd6c variants with Buc condensates. Number of cells analyzed is indicated on the top.

## DISCUSSION

Mechanisms that rely on phase-separation often require the involvement of proteins with multi-valency: the ability to interact with several substrates at the same time. Such multi-valency can stem from multiple folded domains, but also from intrinsically disordered regions (IDRs) or Prion-like domains (PrLDs) (Banani et al., 2016, 2017; Shin & Brangwynne, 2017; Shorter & Lindquist, 2005). Interestingly, germline specification of different species has been characterized by the influence of phase-separated structures, raising the interest towards proteins and domains that regulate these processes in the light of germ cell specification (Boke et al., 2016; Bose et al., 2022; Brangwynne et al., 2009). In zebrafish, two multi-Tudor domain containing proteins, Tdrd6a and Tdrd6c, have appeared as candidates to play a role in the dynamics of zebrafish germplasm, both during oogenesis and embryogenesis. In this discussion we will point out the main implications of our study.

### Tdrd6 proteins do not affect oocyte maturation

To our surprise, Tdrd6c did not enrich within the Bb during oogenesis, unlike its paralog Tdrd6a (Roovers et al., 2018). Instead, the localization of Tdrd6c-mKate2 (Fig. 2B) indicates that Tdrd6c is particularly enriched only within the perinuclear *nuage* of oocytes, while in the rest of the cytoplasm its signal is very weak. Possibly, this difference between Tdrd6a and Tdrd6c stems from the differential interactions of Tdrd6a and Tdrd6c with Buc. In any case, in *tdrd6a; tdrd6c* double mutant females we did not detect major Bb defects, indicating that that the interactions between the Tdrd6 paralogs and Buc are not critical for its function. However, these results do not exclude minor defects in Bb function.

Given that both Tdrd6 paralogs are found in perinuclear *nuage* in oocytes, they could have a role in the regulation of this structure and its function. However, also here, their role is likely modest, as disruption of perinuclear *nuage* in germ cells (by loss of Tdrd1) has been reported to lead to severe germ cell defects in zebrafish (Huang et al., 2011), a phenotype not observed in the *tdrd6* mutants. Indeed, in oocytes mutant for *tdrd6a* and *tdrd6c* perinuclear *nuage* appears to form correctly, as judged from staining of Zili (Fig. S6) (Houwing et al., 2007).

### Tdrd6 proteins are essential for germplasm stability and PGC specification

After fertilization, both Tdrd6a and Tdrd6c are found enriched in the germplasm droplets that aggregate at the cleavage furrows (Fig. 1 and 2). When one of the two proteins is absent in embryos, germplasm defects are not significant in our analyses, which were based on Buc-eGFP protein localization. However, in both scenarios, PGC numbers are reduced at the larval stage, indicating that the germline specification is impaired to some degree during early stages of development (Fig. 3). Possibly, this stems from defects in mRNA recruitment to the germplasm, as Roovers et al. did describe mild, but significant germplasm defects, based on mRNA *in situ* staining (Roovers et al., 2018). Given the association between Tdrd6a (and likely Tdrd6c) and many different mRNPs (Roovers et al., 2018), it is possible indeed that germplasm as assessed by Buc-eGFP versus mRNA *in situ* stainings may lead to different results in absence of Tdrd6 proteins.

Strikingly, in absence of both Tdrd6 proteins, germplasm is lost during the first three hours of development, therefore the embryo is unable to specify PGCs and mature into fertile fish (Fig. 3,4, S3, MovieS2 and MovieS3). While we did not reveal the molecular mechanisms with which Tdrd6a and Tdrd6c contribute to germplasm stability, our previous study on Tdrd6a highlighted how Tdrd6a affects Buc mobility in aggregates, suggesting a direct role in the regulation of the phase-separation properties of germplasm (Roovers et al., 2018). Given the different modes of interaction with Buc, Tdrd6a and Tdrd6c may differentially affect Buc aggregation, or other aspects of germplasm function. Potential differential effects of Tdrd6a and Tdrd6c on Buc aggregation will require studies with purified recombinant proteins. We note that the loss of germplasm over time in absence of Tdrd6 proteins may also be the result of the gradual loss of mRNPs from the germplasm.

### The PrLD of Tdrd6c is prone to aggregate and interacts with Buc

Previous observations showed that Tdrd6a interacts with Buc through di-methylated arginine residues in the Buc C-terminal region (Roovers et al., 2018). Since these are known to be bound by eTD domains, the eTD domains of Tdrd6a, notably the two C-terminal eTDs of Tdrd6a, are likely involved in this interaction. In contrast, when we analyzed Tdrd6c domains in *BmN4* cell, we observed that combinations of Tdrd6c eTDs did not interact with Buc (Fig. 6). On the other hand, we show that the Tdrd6c PrLD forms self-aggregating structures to which Buc droplets can anchor (Fig. 5 and 6). Buc did not merge into these 6c-PrLD structures, but only docked onto them. Furthermore, we found that the first three eTDs of Tdrd6c can modulate the PrLD aggregation behavior, triggering the formation of small cytoplasmic droplets, instead of the large, irregular aggregates formed by the PrLD alone. Additionally, Buc merged with these granules, instead of docking onto them (Fig. 6). Interestingly, the first three eTDs of Tdrd6c are missing the aromatic cage and the key asparagine residue responsible for interaction with methylated substrates (Fig. S2, S3) (H. Liu et al., 2010; K. Liu et al., 2010). This would suggest that the first three eTDs may have affinity for non-methylated residues within the Tdrd6c PrLD, thereby modulating its aggregation behavior. Such non-methylation dependent interaction between an eTD and a substrate has been shown before (Webster et al., 2015).

Altogether, the differences in Buc interactions between Tdrd6a and Tdrd6c are intriguing, since *in vivo*, they act (at least partially) redundantly. That said, we cannot exclude that the differences we describe hold a key to biological functionalities that did not become evident in our current studies. It may well be that the germplasm, and/or the germ cells formed in *tdrd6* single mutants do display defects. However, these would have to be compatible with germ cell function.

## MATERIAL AND METHODS

### Zebrafish lines

Zebrafish strains were housed at the Institute of Molecular Biology in Mainz and bred and maintained under standard conditions (28°C water temperature and lighting conditions in cycles of 14:10 hours light:dark) (Westerfield, 1995. Larvae < 5 days post fertilization were kept in E3 medium (5 mM NaCl, 0.17 mM KCl, 0.33 mM CaCl2, 0.33 mM MgSO4) at 28°C. The zebrafish strains used for this work are listed in the table below (Table 1). Periodic out-crossing of each line was done with mating to zebrafish wild type (wt) strains (AB or TUE). All experiments were conducted according to the European animal welfare law and approved and licensed by the ministry of Rhineland-Palatinate.

**Table 1.** List of zebrafish alleles. Names, descriptions and references of the zebrafish alleles used for this work.

| Allele | Description | References |
| --- | --- | --- |
| *tdrd6c<sup>mz69</sup>* | CRISPR-induced knock out mutant allele | This paper |
| *tdrd6a<sup>Q185X</sup>* | Knock out allele from ENU mutagenized library | (Roovers et al., 2018; Wienholds et al., 2002) |
| *tdrd6c<sup>mz87tg</sup>* | Transgenic line expressing Tdrd6c – mKate2 protein | This paper |
| *Tg(vasa:eGFP)* | Transgenic line expressing eGFP protein from vasa promoter | (Krøvel & Olsen, 2002) |
| *Tg(buc:buc-eGFP)* | Transgenic line expressing Buc-eGFP protein | (Riemer et al., 2015) |

### Cell culture

*BmN4* cells were cultured at 27°C in IPL-41 (Gibco) medium supplemented with 10% Fetal Bovine Serum (Gibco) and 0.5% Pen-Strep. For imaging, cells were grown in 8-well µ-slides (Ibidi, Cat# 80826).

### Design and cloning of CRISPR-guide-RNAs

Crispr-guide RNAs (gRNAs) were designed using CRISPRscan in the UCSC genome browser (Moreno-Mateos et al., 2015). Designed oligonucleotides were produced by Integrated DNA Technology and incorporated into pDR274 plasmid via restriction enzyme digestion followed by T4 ligation (NEB, Cat#M0202) (Hwang et al., 2013).

### Expression of CRISPR-guide-RNAs

Plasmids containing gRNAs encoding sequences were linearized with DraI restriction enzyme (NEB, Cat# R0129L) and extracted from gel (QIAquick Gel Extraction Kit, Cat# 28704). Linearized sequences were used for RNA *in vitro* transcription (MEGAscript SP6 Transcription Kit, Cat# AM1330). Subsequently, expressed gRNAs were treated with Turbo DNase (Invitrogen, Cat# AM2239) for 30 min. at 37°C and purified by precipitation and re-suspended in RNase free water.

### RNA extraction and cDNA preparation

Ovaries from fish were homogenized in TRIzol Reagent (Thermo Fisher, Cat# 15596018) using a sterile pestle in a 1.5 mL sterile tubes. Samples were centrifuged at 12000 g, at 4°C, for 5 min. and the clear supernatant was transferred to a new tube. To each tube, 0.2 mL of chloroform were added. Tubes were vortexed for 20-30 seconds and then incubated for 2-3 min. Samples were centrifuged for 15 min. at 4°C at 12000 g. The upper aqueous phase was transferred to new tubes and 0.5 mL of isopropanol were added. Samples were incubated 1 hour at −80°C and then centrifuged for 10 minutes at 4°C at 16000 g. Supernatant was discarded and pellet re-suspended in 1 mL of 75% Ethanol (EtOH). Samples were vortexed and centrifuged for 5 min. at 160000 g at 4°C. Supernatant was discarded and samples were let dry. Pellets were re-suspended in 20 µL of sterile water.

### Gateway Cloning of *tdrd6c-mKate2*

The plasmid containing the *tg(ziwi:tdrd6c-mKate2-tdrd6c3’UTR)* sequence was generated with the Tol2kit for multisite-Gateway cloning (Kwan et al., 2007). The *tdrd6c* ORF and the *tdrd6c-3’utr* were amplified from zebrafish ovarian cDNA. The *mKate2* sequence was amplified from existing plasmid in the lab and annealed to the *tdrd6c-3’utr* with incubation at 95°C for 5 min. in a Thermocycler and gradual cool down. The *tdrd6c* ORF sequence and the *mKate2-tdrd6c-3’utr* sequence were cloned respectively into the pDonr221 and the p2r-P3 vectors using BP clonase reactions (Thermo Fisher, Cat# 11789020). The LR reaction (Thermo Fisher, Cat# 12538120) was performed using the two obtained plasmids from the BP reaction and the p5E_pziwi (containing the *ziwi promoter*) together with the destination vector tol2CG2 (Kwan et al., 2007).

### Injection of zebrafish zygotes

CRISPR-gRNAs were diluted in 0.05% Phenol red (Sigma-Aldrich, Cat# 143-74-8) to a final concentration between 100-500 ng/µL, together with 2 µM of Cas9 protein (NEB, Cat# M0646T). Tol2-plasmids were diluted in 0.05% Phenol red to a final concentration of 100 ng/µL, together with Transposase encoding mRNA (100 ng/µL). A volume between 2 and 5 nL (1 to 2.5 µg of each RNA or 200 ng to 500 ng of plasmid) was injected to wild type zebrafish zygotes via glass needles capillaries (Harvard Apparatus, Cat# EC1 30-0038).

### Genotyping of zebrafish strains

DNA was extracted from caudal fin tissue amputated from anesthetized fish. Fins were incubated in Lysis Buffer (50 mM KCl, 2.5 mM MgCl2, 10 mM Tris, 0.5% NP40, 0.5% Tween20, 0,01% w/v gelatine and 0,1 mg/mL Proteinase K, pH 8) for 1 hour at 60°C, followed by proteinase K inactivation for 15 min. at 95°C.

### Gibson Assembly Cloning

Plasmids used for transfection of *BmN4* cells and for protein expression were cloned using Gibson Assembly strategy (Gibson et al., 2009), using the reagents produced by the IMB Protein Production Core Facility (Gibson Assembly Reaction Mix) and following their provided protocols.

### PGC counts

At 24 hpf, eGFP-positive zebrafish embryos, obtained from vasa:eGFP positive mothers, were dechorionated and fixed in 4%PFA/PBS-Tw. Then, samples were washed twice in PBS and placed in 8-wells slides (Ibidi, Cat# 80826) and analyzed under the 10X objective of the Visitron VisiScope microscope and GFP-positive cells were counted manually.

### Anti-Tdrd6c antibody production

With the help of the IMB Protein Production Core Facility, I selected, cloned and expressed the following Tdrd6c-peptide:

H2N-LPEPCELNEWFKNYATDCMFNVVVKLNSSGKLSVEMYDDKTNLNLKIKDLWSKTK-CONH2.

This was fused N-terminally with a His-MBP tag to favor solubility and purification. The tag could be cleaved thanks to a 3C Protease-recognition site.

Expression was performed in BL21DE3 competent cells (Table 2). Cells were lysed with Lysis Buffer (50 mM Tris, 300 mM NaCl, pH 7) and the uncleaved peptide was purified with IMAC and eluted in Elution Buffer (50 mM Tris, 300 mM NaCl, 350 mM Imidazole, pH 7). His-MBP tags were cleaved off o/n with 3C Protease. The cleaved peptide was purified with reverse IMAC and gel filtration.

**Table 2.** List of cellular bacterial strains used in this Thesis. Names, description of their usage and identifiers of bacterial strains used for different cloning strategies and purposes of this work.

| Name | Usage within this Thesis | Identifier |
| --- | --- | --- |
| Subcloning Efficiency™<br>DH5α™ Competent Cells | General cloning strategies | Thermo Fisher<br>(Cat# 18265017) |
| One Shot™ Top10<br>Chemically Competent<br>Cells | Cloning of Tdrd6c-epitope | Thermo Fisher<br>(Cat# C404003) |
| BL21(DE3)<br>Competent Cells | For Tdrd6c-epitope expression | Thermo Fisher<br>(Cat# EC0114) |
| One Shot™ *ccdB*<br>Survival™<br>2 T1<sup>R</sup> Competent Cells | Cloning of *tdrd6c-mKate2* final<br>plasmid | Thermo Fisher<br>(Cat# A10460) |

Tdrd6c antibodies were raised in rabbits with the purified cleaved peptide (Eurogentec). Antisera were subsequently purified against the cleaved peptide by the IMB Protein Production Core Facility.

### Whole mount Immuno-Histochemistry

Zebrafish samples (ovaries, embryos or larvae) stored in MetOH at −20°C were rehydrated in PBS-Tween/MetOH series (50%, 75%, 87.5% and 100%). Two additional washing steps in 1% Triton X-100 in PBS were used for ovaries and larvae. Antigen-retrieval was performed with incubation of the samples in 150 mM Tris (pH 9) solution at RT for 5 min. and then at 70°C for 15 min. Samples were then blocked in Blocking Solution (8% BSA in PBS-Tw) for 90 min at RT with gentle agitation. After blocking, samples were incubated o/n at 4°C with dilution of primary antibodies (anti-Tdrd6a 1:500, anti-Tdrd6c 1:100, anti-Zili 1:500 and anti-Ziwi 1:100) in Blocking Solution and with gentle agitation. Samples were washed 6 times (30 min. each wash) in PBS-Tw and then incubated in Blocking Solution with 1:500 dilution of secondary antibodies. Samples were washed in PBS and gently transferred to Ibidi slides (8-wells slides for embryos and larvae, 2-wells slides for ovaries).

### Live imaging of Zebrafish embryos

Zebrafish embryos were collected in petri dishes containing E3 solution manually de-chorionated. With glass pipettes, embryos at the 1-cell stage were gently transferred to 8-wells slides (Ibidi, Cat# 80826). Gently, E3 in excess was removed keeping the samples under the level of the surface of the solution and PBS containing ∼0.5% Low Gelling agarose was added one drop at the time. Samples were imaged in the VisiScope system (Leica) with 20X objective.

### BmN4 cell transfection and imaging

*BmN4* cells grown in 25 mL flasks were re-suspended and counted automatically (Bio-Rad, TC20 Automated Cell Counter). Circa 80 thousand cells were seeded in each well of an 8-wells-slides (Ibidi, Cat# 80826) in a volume of 300 µL of IPL-41 medium (Gibco) supplemented with Fetal Bovine Serum (Gibco) and 0.5% Pen-Strep. The next day, 1 µL of each plasmid (from stock of 100 ng/µL) was in solution with 0.5 µl of X-tremeGENE_HP (Roche) and 28.5 µL of IPL-41 (Gibco) medium supplemented with 10% Fetal Bovine Serum (Gibco) for 20-30 min. prior to transfection. Transfected cells were imaged 2 days after transfection at the Leica TCS SP5 Confocal microscope with 63X objective.

### Quantification and Statistical Analysis

Quantifications of microscopy images were carried out either manually or with pipelines within the Arivis Sofware (ZEISS).

A two-sided Wilcoxon test was used to calculate significant differences between populations of maternal mutants in the following experiments: measurement of PGC and male abundancy. Tests were corrected for multiple testing within the respective data set.

For the statistical analysis of germplasm condensate numbers, a quasi-Poisson model was used. Differences between the *control* group and the other groups were assessed with Wald-tests on the model parameters.

For the statistical analysis of germplasm condensate volumes, a Gaussian linear mixed effects models (LMM) were used. Differences between the *control* group and the other groups were assessed with t-tests on the model parameters(Bates et al., 2015).

## Tables of Key Resources

**Table 3.** List of chemicals and kits used in this Thesis.

| **Reagent** | **Source** | **Identifier** |
| --- | --- | --- |
| **Antibodies** |  |  |
| anti-Tdrd6c (from Rabbit) | This Thesis | Epitope:<br>LPEPCELNEWFKNYATDCM<br>FNVVVKLNSSGKLSVEMYD<br>DKTNLNLKIKDLWSKTK |
| anti-Tdrd6a (from Rabbit) | (Roovers et al., 2018) | Epitope: QAVVHEPESEKEKRD |
| anti-Zili (from Rabbit) | (Houwing et al., 2007) | N/A |
| anti-Ziwi (from Rat) | (Roovers et al., 2018) | Epitope: QLVGRGRQKPAPGAM |
| anti-Rabbit-Alexa555 | Invitrogen | Cat# A21428 |
| anti-Rabbit-Alexa647 | Abcam | Cat# ab150075 |
| anti-Rat-Alexa555 | Invitrogen | Cat# A21434 |
| anti-Rat-Alexa647 | Abcam | Cat# ab150155 |
| **Chemicals** |  |  |
| TRIzol | Thermo Fisher | Cat# 15596018 |
| TRIzol LS | Thermo Fisher | Cat# 10296010 |
| Ethyl 3-aminobenzoate<br>methanesulfonate (E10521) | Sigma-Aldrich | Cat# 886-86-2 |
| 4%-12% NuPage NOVEX<br>gradient gel | Thermo Fisher | Cat# NP0321 |
| NuPAGE LDS sample (Buffer<br>4x) | Thermo Fisher | Cat# NP0007 |
| Paraformaldehyde 20% | Sigma-Aldrich | Cat# 252549 |
| PBS | From IMB Media Lab<br>Core Facility | - |
| Bovine Serum Albumine (BSA) | Sigma-Aldrich | Cat# A7906 |
| Dextran Sulfate | Sigma-Aldrich | Cat# 42867-5G |
| Vanadyl-ribonucleoside<br>complex | NEB | Cat# S1402S |
| ProLong_Gold Antifade<br>Mountant | Thermo Fisher | Cat# P10144 |
| Triton X-100 | Sigma-Aldrich | Cat# T9284 |
| Tween20 | Sigma-Aldrich | Cat# P1379 |
| cOmplete Mini, EDTA-free<br>protease inhibitor<br>cocktail Tablets | Roche | Cat# 11836170001 |
| Ficol PM 400 | Sigma-Aldrich | Cat# 26873-85-8 |
| IPL-41 insect medium | Gibco | Cat# 11405057 |
| 8-well m-slides | Ibidi | Cat# 80826 |
| 9 mL X-tremeGENE_ HP | Roche | Cat# 6365779001 |
| BP clonase II | Thermo Fisher | Cat# 11789020 |
| LR clonase II plus | Thermo Fisher | Cat# 12538120 |
| **Critical Commercial Assays** |  |  |
| Sp6 mMESSAGE MACHINE kit | Invitrogen | Cat# AM1340 |
| M-MLV reverse transcriptase,<br>RNase H<br>point mutant | Promega | Cat# M3681 |
| QIAquick PCR Purification Kit | Qiagen | Cat# 28106 |

**Table 4.** List of Online resources used in this Thesis.

| Name | URL | Reference |
| --- | --- | --- |
| Ensembl | http://www.ensembl.org/index.html | (Martin et al., 2023) |
| CrisprSCAN | https://www.crisprscan.org/ | (Moreno-Mateos et al., 2015) |
| IDT Oligo | https://eu.idtdna.com/pages/products/custom-dna-rna/dna-oligos/custom-dna-oligos | - |
| PLAAC | http://plaac.wi.mit.edu/ | (Lancaster et al., 2014) |
| IUPred2 | https://iupred2a.elte.hu/ | (Dosztanyi et al., 2005;<br>Dosztányi, 2018;<br>Mészáros et al., 2018) |
| ClustalOmega | https://www.ebi.ac.uk/Tools/msa/clustalo/ | - |
| NCBI | https://www.ncbi.nlm.nih.gov/Structure/cdd/wrpsb.cgi | (Lu et al., 2020;<br>Marchler-Bauer et al.,<br>2011, 2015, 2017;<br>Marchler-Bauer &<br>Bryant, 2004) |
| StarSEQ | https://www.starseq.com/life-science/sanger-sequencing/u-mix/ | - |

## ACKNOWLEDGMENTS

We thank the members of our laboratory for fruitful discussions. We thank the following IMB Core Facilities for their contribution and valuable support: Microscopy, Protein Production and the Media lab. In particular, we thank Martin Möckel, for the assistance in the design and purification of the Tdrd6c epitope and the purification of the resulting antibody. The work was supported by core funds from IMB to RFK.

---

**Fig. S1.**
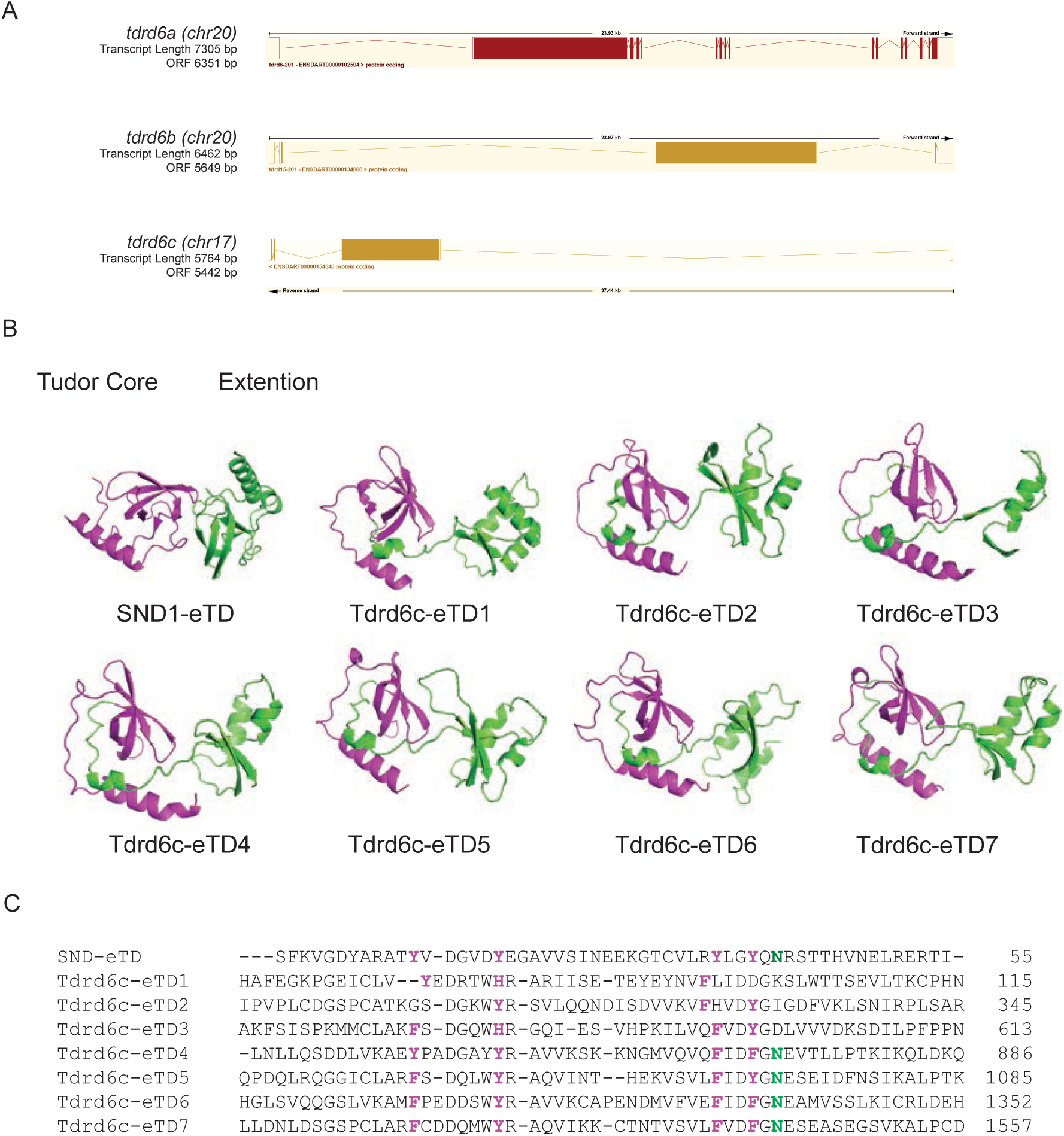
TDs of Tdrd6c are extended TDs. (A) View of the zebrafish *tdrd6 loci* as annotated from Ensembl database. (B) AlphaFold predictions of the 7 eTDs of Tdrd6c compared to the eTD of the SND-1 protein; in magenta the Tudor core and in green the extended domain. (C) Alignment of the amino-acid sequences of the 7 eTDs of Tdrd6c and the eTD of SND-1; highlighted: in magenta are residues contributing to the aromatic-cage of canonical eTDs; in green the Asparagine residue that also interacts with methylated-Arginines of a substrate.

**Fig. S2.**
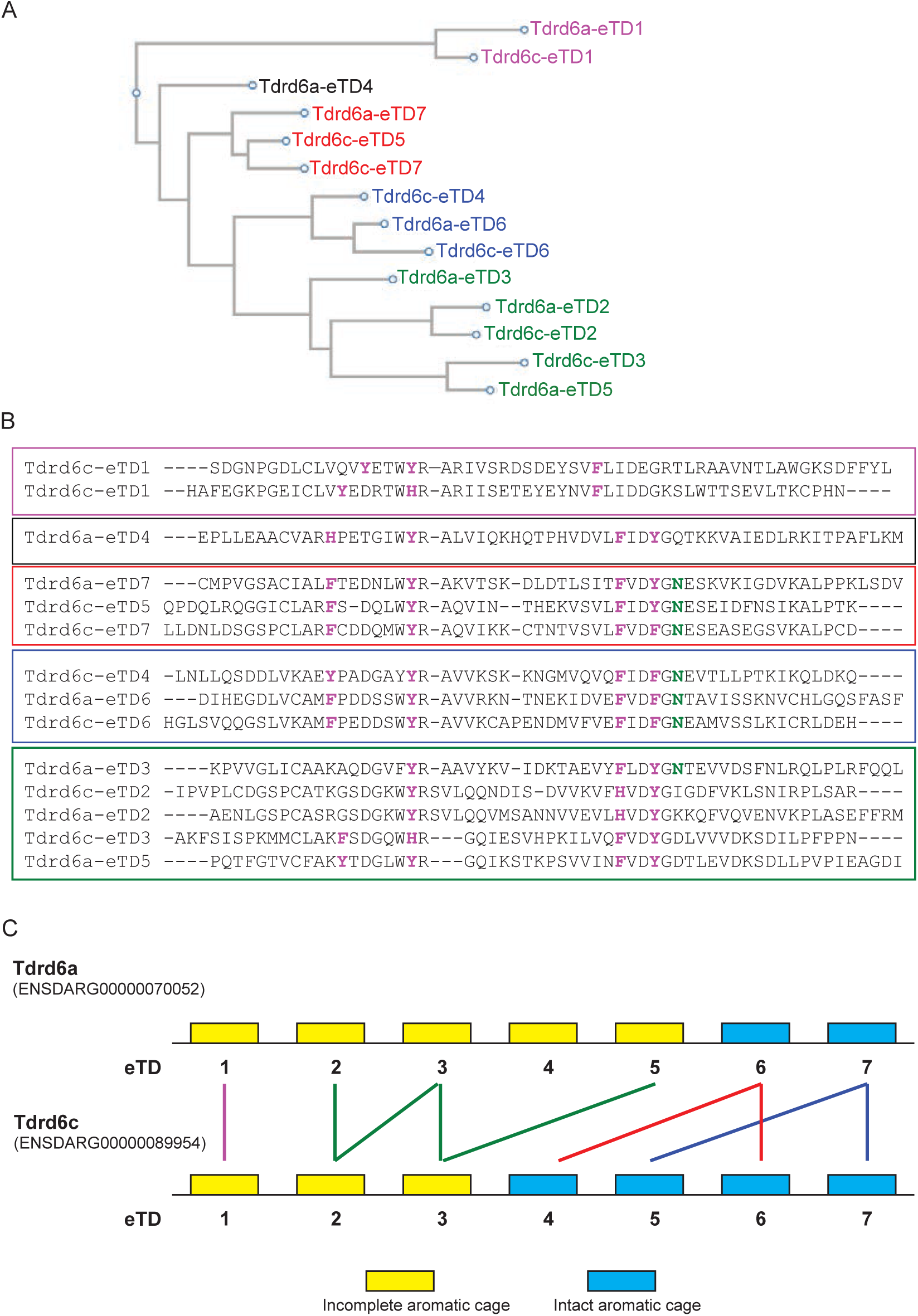
Comparison of eTDs of Tdrd6a and Tdrd6c. (A) Cladogram of eTDs from Tdrd6a and Tdrd6c, highlighting 5 groups (color coded) of eTDs. (B) Sequence alignment of the eTDs of Tdrd6a and Tdrd6c, used to generate panel A in CLUSTALW. Colored boxes, matching panel A, highlight the different groups. Residues in red are the ones supposed to contribute to the formation of the aromatic cage, in green the Asparagine residue that should bind methylated-arginine residues. (C) Schematic representation of Tdrd6a and Tdrd6c protein structure with their eTDs. Colored lines refer to the cladogram and indicate similarity between different eTDs.

**Fig. S3.**
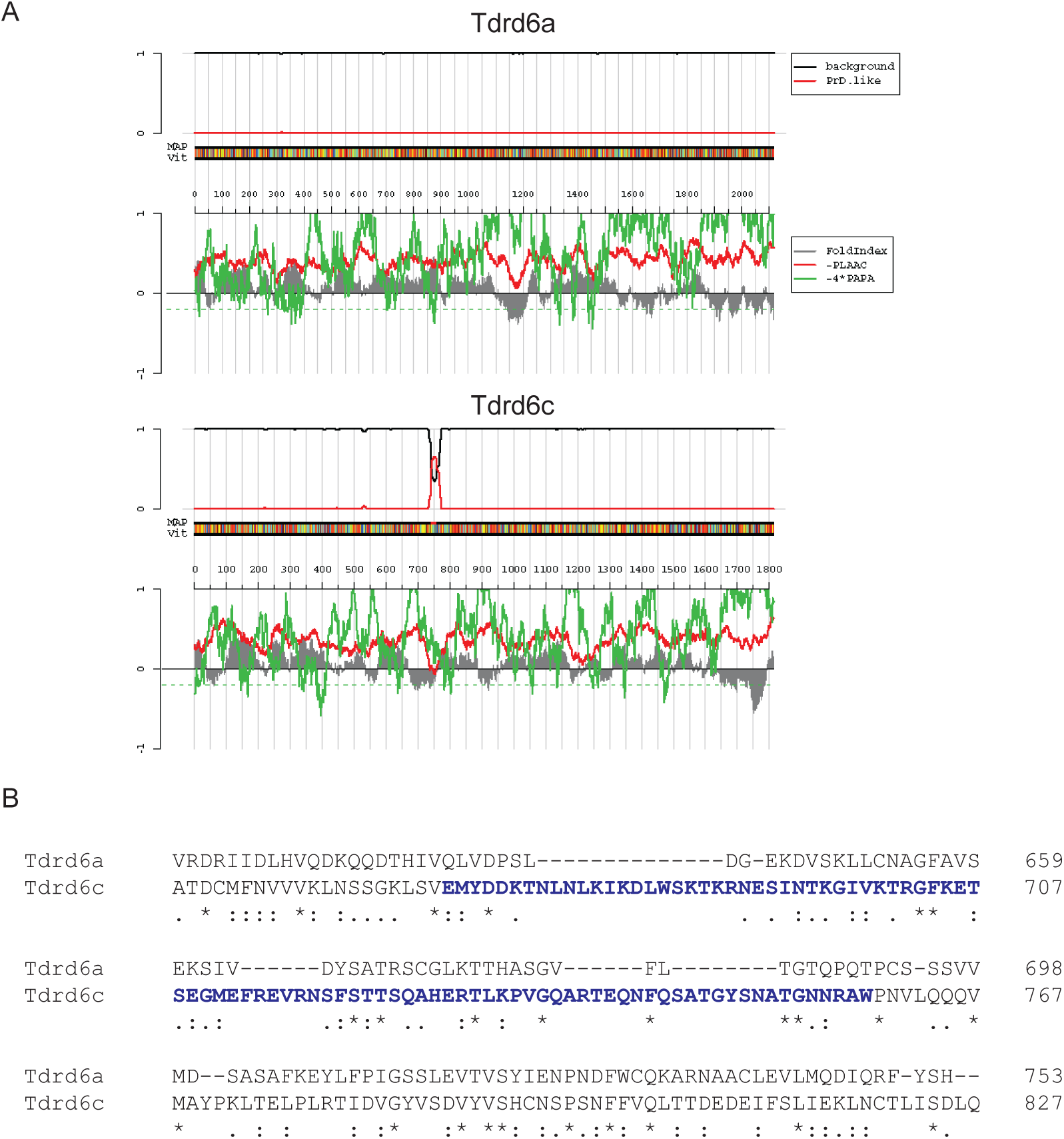
Tdrd6c predicted Prion-like domain. (A) PLAAC prediction of Prion-like domains in Tdrd6a (Top) and in Tdrd6c (Bottom) (Lancaster et al., 2014). (B) Alignment of a part of Tdrd6a and Tdrd6c highlighting, in blue, that Tdrd6c predicted Prion-like domain is a unique sequence between the 2 proteins.

**Fig. S4.**
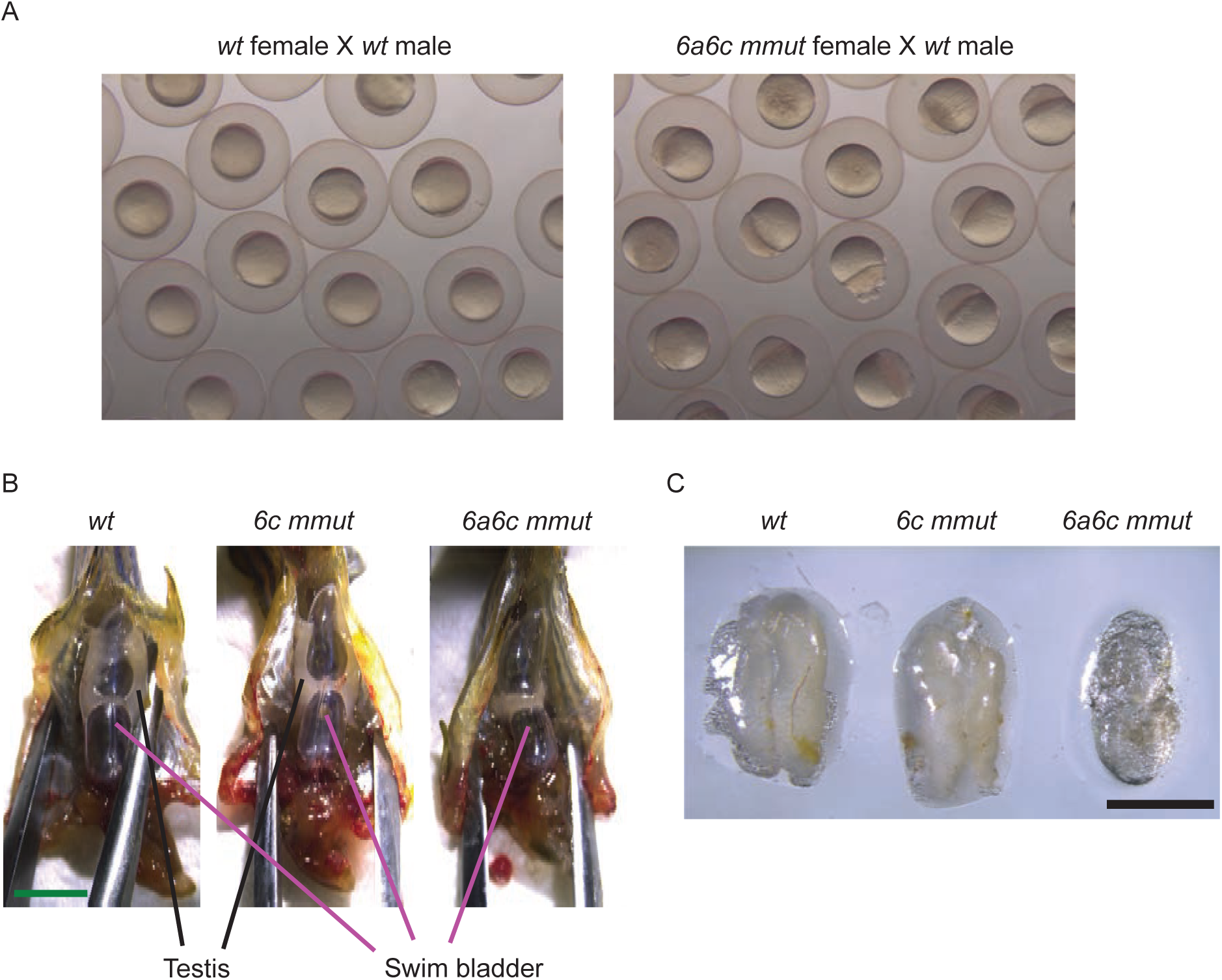
Maternal Tdrd6a and Tdrd6c are required for normal gonad morphology. (A) Collection of offspring from wt fathers (left) and from *6a6c mmut* fathers (right); wt eggs are fertilized and develop correctly when mating occurs with *wt* fathers while they appear unfertilized when collected from mating of *6a6c mmut* males. (B) Dissection of *6a6c mmut* zebrafish reveals that their gonads are not developed, in contrast to *wt* and *6c mmut* fish (scale bar = 5 mm). Only a small rudimentary part of the gonad structure can be seen crossing the swimming bladder. (C) Isolation of gonads from *wt, 6c mmut* and *6a6c mmut* further highlights that gonads fail to mature in *6a6c mmut* (scale bar = 3 mm). In wt and *6c mmut* samples the fat patch was overlain by milky-white tissue, which is the testis. In the *6a6a mmut* sample only the fat patch was visible, and testis tissue could not be clearly distinguished.

**Fig. S5.**
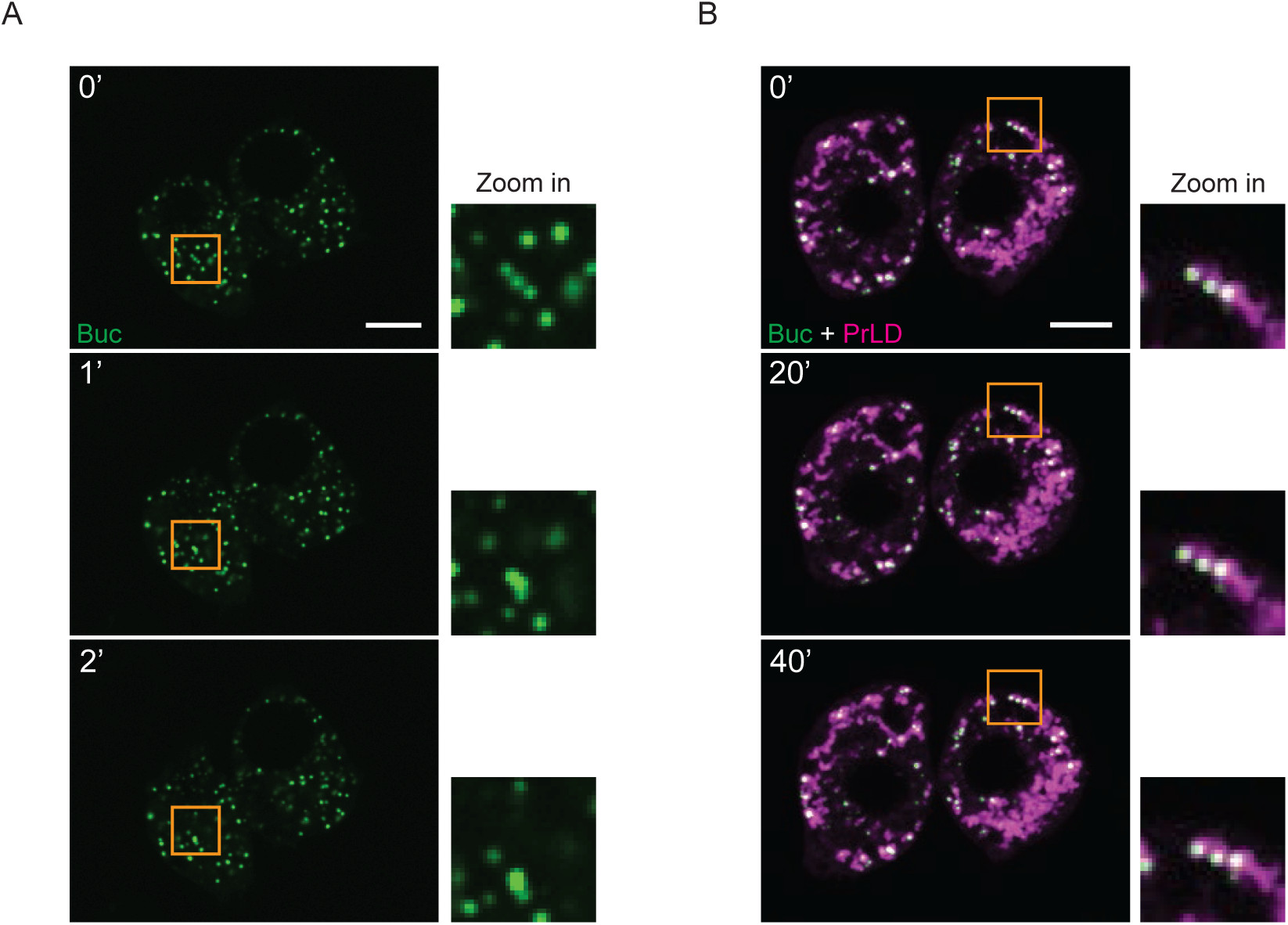
BucGFP mobility in BmN4 cells. (A) Buc-eGFP expression observed in BmN4 cells over 2 minutes. The displayed images are still from Movie S4. (B) Buc-eGFP and mCherry-PrLD co-expression observed over 40 minutes in BmN4 cells. The displayed images are still from Movie S5. (C) Orange boxes indicate areas of zoom in. Scale bars = 10 µm.

**Fig. S6.**
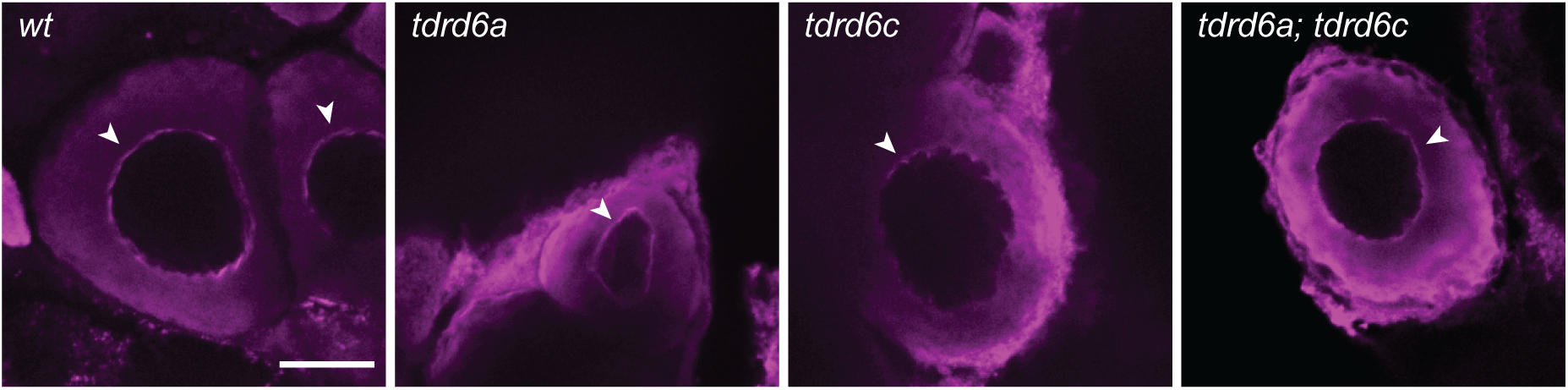
The *nuage* appears correctly formed in *tdrd6a* and *tdrd6c* mutant oocytes. Perinuclear *nuage* (magenta, arrowheads) stained with antibody against Zili in *wt* and mutant oocytes. Scale bar = 50 µm.

**MovieS1. Localization of Tdrd6c-mKate2 during embryogenesis.** Live imaging displaying the localization of Tdrd6c-mKate2 (magenta) throughout the first 3 hours of development. In green Buc-eGFP marking the germplasm.

**MovieS2. Distribution of germplasm in *wt embryos*.** Germplasm visualized via Buc-eGFP (green) in *wt* embryos over the first 3 hours of development.

**MovieS3. Distribution of germplasm in *6a6c mmut embryos*.** Germplasm visualized via Buc-eGFP (green) in *6a6c mmut* embryos over the first 3 hours of development.

**MovieS4. BucGFP behavior in BmN4 cells.** Live imaging of Buc-GFP (green) expressed in BmN4 cells (time-lapse of 3 minutes). Scale bar = 10 µm.

**MovieS5. BucGFP behavior in BmN4 cells co-expressing mCherry-PrLD.** Live imaging of Buc-eGFP (green) expressed in BmN4 cells together with mCherry-PrLD (magenta) (time-lapse of 12 hours). Scale bar = 10 µm.

